# DOCK2 sets the threshold for entry into the virtual memory CD8^+^ T cell compartment by negatively regulating tonic TCR triggering

**DOI:** 10.1101/582486

**Authors:** Ezana Demissie, Vinay S Mahajan, Faisal Alsufyani, Sudha Kumari, Grace J Yuen, Vinayak Viswanadham, Johnson Q. Tran, James J. Moon, Darrell J Irvine, Shiv Pillai

## Abstract

The control of cytoskeletal dynamics by Dedicator of cytokinesis 2 (DOCK2), a hematopoietic cell-specific actin effector protein, has been implicated in TCR signaling and T cell migration. Biallelic mutations in *Dock2* have been identified in patients with a recessive form of combined immunodeficiency with defects in T, B and NK cell activation. Surprisingly, we show here that certain immune functions of CD8^+^ T cells are enhanced in the absence of DOCK2. *Dock2*-deficient mice have a pronounced expansion of their memory T cell compartment. Bone marrow chimera and adoptive transfer studies indicate that these memory T cells develop in a cell-intrinsic manner following thymic egress. Transcriptional profiling, TCR repertoire analyses and cell surface marker expression indicate that *Dock2*-deficient naive CD8^+^ T cells directly convert into virtual memory cells without clonal effector T cell expansion. This direct conversion to memory is associated with a selective increase in TCR sensitivity to self-peptide MHC *in vivo* and an enhanced response to weak agonist peptides *ex vivo*. In contrast, the response to strong agonist peptides remains unaltered in *Dock2*-deficient T cells. Collectively, these findings suggest that the regulation of the actin dynamics by DOCK2 enhances the threshold for entry into the virtual memory compartment by negatively regulating tonic TCR triggering in response to weak agonists.

## INTRODUCTION

CD8^+^ T cells are critical effectors of cellular adaptive immunity. A cardinal feature of T cell responses is the establishment of long-lived memory cells that rapidly respond upon pathogen re-exposure. It is widely held that cognate peptide activation drives naive T cell clonal expansion and subsequent contraction, leaving behind memory cells. Antigen-induced memory cells persist following the clearance of an acute infection (Kaech et al., 2003; Yu et al., 2017; Akondy et al., 2017). CD8^+^ T cell memory cell formation that is not accompanied by effector cell differentiation, but that shares most of the transcriptional and functional traits of conventional memory, has also been extensively documented (Jameson, 2002; Goldrath et al., 2004; Hamilton and Jameson, 2008; Cheung et al., 2009). Memory T cells that are apparently cognate antigen-independent are also found in the sterile environment of the human fetal spleen and cord blood (Zhang et al., 2014; Jacomet et al., 2015; Van Kaer, 2015), suggesting that the generation of these cells is the product of a physiologically relevant T cell differentiation pathway. These cells have also been seen in germ-free mice, consistent with their dependence on homeostatic signals and self-peptides for generation and survival, as opposed to triggering by microbial antigens. Fate-mapping of murine CD8^+^ T cells based on their developmental timing has shown that CD8^+^ T cells that develop early in life have a greater propensity to differentiate into virtual memory cells (Smith et al., 2018).

The phenomenon of “virtual” memory T cells was initially observed when naive T cells were adoptively transferred into lymphopenic settings such as neonatal and irradiated mice (Goldrath and Bevan, 1999; Ernst et al., 1999; Min et al., 2003), and was termed lymphopenia-induced proliferation. Lymphopenia-induced proliferation requires a combination of common γ-chain cytokines such as IL-7 or IL-15 and TCR triggering by self-peptides (Tan et al., 2001). A transient increase in tonic “survival” signals may be sufficient to drive such memory cell differentiation (Sprent and Surh, 2011). Consistent with this idea, there is a direct correlation between lymphopenia-induced conversion to memory and the strength of tonic self-peptide signaling that a T cell receives (Ge et al., 2001; Kieper et al., 2004).

Several studies have documented the presence of cognate antigen-independent CD8^+^ T cell memory that arises without any apparent lymphopenia in naive lymphoreplete mice (White et al., 2017; Haluszczak et al., 2009). These memory-phenotype cells arise in the periphery, requiring common γ-chain cytokines (mainly IL-15) (Sosinowski et al., 2013; Fiege et al., 2015) and self-peptide TCR triggering (White et al., 2016), in a manner reminiscent of lymphopenia-induced memory cells. Studies in mice lacking the IL-4R showed that the production of CD8^+^ virtual memory cells in the periphery is enhanced by IL-4 (Akue et al., 2012). CD8.4 mice, expressing CD8 fused to a CD4-derived intracellular domain, exhibit higher levels of CD8-Lck coupling and have an expanded CD8^+^ virtual memory compartment, suggesting that the intrinsic sensitivity of the TCR signaling machinery can regulate the size of the CD8^+^ virtual memory compartment (Drobek et al., 2018). Virtual memory cells also provide antigen-independent innate-like bystander protection in the context of intracellular infection (White et al., 2016), and contribute to protective immunity in mice (Berg et al., 2003; Chu et al., 2013) and humans (Kim et al., 2017).

Aside from homeostatic cytokine signaling and tonic TCR triggering, very little is known about the regulators of the CD8^+^ T cell virtual memory compartment. FYB1 (Fyn binding protein 1) has been proposed to function as a negative regulator of the size of the CD8^+^ virtual memory compartment by limiting the response to IL-15 (Fiege et al., 2015). In this study, we used mice lacking DOCK2, a hematopoietic-restricted guanine exchange factor (GEF), to show that it functions as a novel negative regulator of the CD8^+^ virtual memory compartment. In some circumstances, CD8^+^ memory-like T cells can also be generated in a non-cell-autonomous manner upon exposure to PLZF^+^ NKT cell-derived IL-4 during thymic development (Lee et al., 2011). To distinguish them from virtual memory T cells that arise in the periphery, such cells have been referred to as “innate” CD8^+^ cells and have been observed in some knockout mouse strains (*e.g*., mice deficient in KLF2, ITK, or CBP), as well as in unmanipulated mice from the BALB/c background, which exhibits a higher baseline level of IL-4 producing iNKT cells (Lee et al., 2011). However, this phenomenon has not been observed in the B6 background, which exhibits a low level of iNKT cells relative to other inbred mouse strains (Lee et al., 2011). Furthermore, the *Dock2*-deficient mice (C57BL/6NHsd background) described in this study have reduced numbers of iNKT cells compared to wild-type C57BL/6J controls (Mahajan et al., 2016).

DOCK2 (Dedicator of cytokinesis 2) activates the actin effector Rho GTPase Rac by catalyzing the transition from the inactive GDP-bound state to the active GTP-bound state (Fukui et al., 2001). DOCK2 localizes to the cell membrane via its DHR1 domain mediated interactions with PIP3 and polybasic amino acid cluster based interactions with phosphatidic acid, thus ensures spatially controlled activation of GTPase Rac at the plasma membrane (Fukui et al., 2001; Nishikimi et al., 2009, 2013). GTP bound RAC1 subsequently drives actin polymerization enabling cytoskeletal rearrangements required for lymphocyte chemotaxis (Fukui et al., 2001; Terasawa et al., 2012; Gotoh et al., 2008), T cell interstitial motility (Nombela-Arrieta et al., 2007), plasmacytoid dendritic cell cytokine secretion (Gotoh et al., 2010), and TCR activation (Sanui et al., 2003; Le Floc’h et al., 2013).

We have previously described a loss-of-function *Dock2* allele (*Dock2^hsd^*) that had been inadvertently introduced into multiple mouse lines (Mahajan et al., 2016). Using this knockout allele of *Dock2* extensively in this study, we find that the responsiveness of *Dock2*-deficient CD8^+^ T cells to weak agonists is unexpectedly enhanced. Thus, while DOCK2 may promote TCR responses to strong agonists, it appears to constrain the responsiveness to weak TCR agonists. *In vivo*, the loss of DOCK2 results in an enhanced conversion to virtual memory T cells. In this present study, we show that the memory phenotype T cells expanded in the absence of DOCK2 are polyclonal virtual memory cells and demonstrate that they are generated by direct conversion of naive T cells into memory as a result of cell-intrinsic hyperresponsiveness of *Dock2^hsd/hsd^* T cells to weak agonists. Mice with other engineered *Dock2* mutations also exhibit the same phenotype. These findings suggest that the absence of DOCK2 lowers the threshold of self-peptide triggering required to enter the virtual memory T cell compartment.

## RESULTS

### Absence of DOCK2 results in the expansion of virtual memory T cells

In an earlier study, we identified an approximately 3-fold expansion of memory phenotype (MP) CD8^+^ T cells in mice carrying a spontaneous loss of function mutation in the guanine exchange factor DOCK2 (*Dock2^hsd/hsd^*) (Mahajan et al., 2016). This expansion is not specific to this particular allele of *Dock2*, as it is also present in gene targeted *Dock2*-deficient mice (Fukui et al., 2001) (Figure 1A). The increase in the percentage of *Dock2^hsd/hsd^* MP cells is also mirrored by a similar increase in the total numbers of these cells (Figure 1B). Notably, these cells lack surface expression of NK1.1 and express both CD8α and CD8β (data not shown).

**Figure 1:**
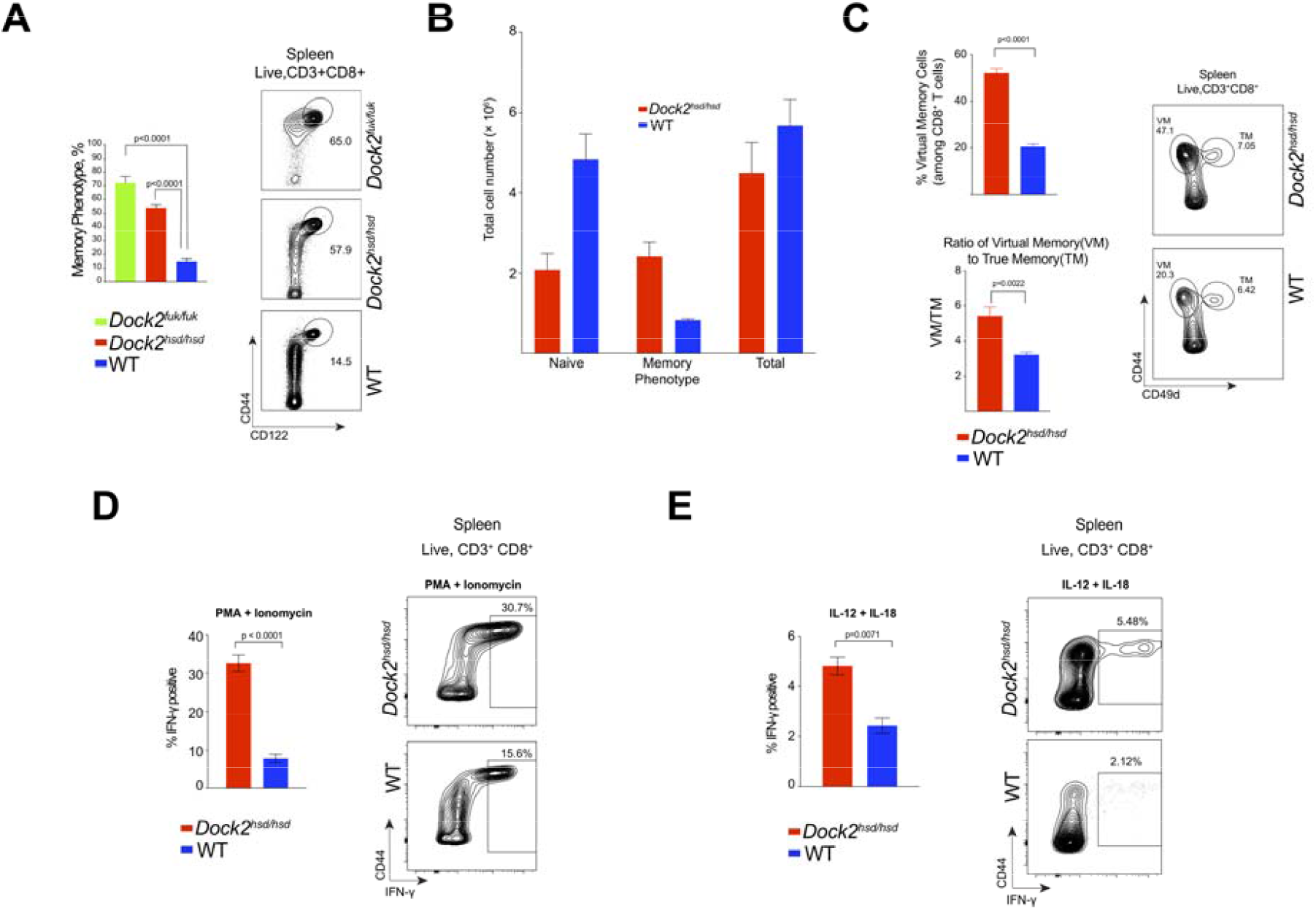
*Dock^hsd/hsd^* mice have an expanded virtual T cell compartment and are resistant to intracellular bacterial infection. (A) The proportion of CD8^+^ memory phenotype cells (MP) in the spleens of *Dock2^hsd/hsd^, Dock2^Fukui/Fukui^* and wild-type (WT) mice as measured by flow cytometry. (B) Number of naive (CD44^lo^CD122^−^), memory phenotype (CD44^hi^CD122^hi^) and total CD8^+^ T cells in the spleens of WT and *Dock2*-deficient mice. (C) Percentage of CD8^+^ virtual memory (VM) (CD44^hi^CD49d^lo^) T cells among total CD8^+^ T cells as well as the ratio of VM to true memory (TM) (CD44^hi^CD49d^hi^) CD8^+^ T cells in the spleens of WT and *Dock2*-deficient mice (D & E) Intracellular staining for IFN-γ in WT and *Dock2*-deficient CD8^+^ T cells stimulated for 4 hours with PMA and Ionomycin (D) or for 18 hours with IL-12 and IL-18 (E).

Recent studies have identified cognate antigen-independent memory cells that arise in unmanipulated lymphoreplete mice. (White et al., 2017; Haluszczak et al., 2009; Akue et al., 2012). Such “virtual” memory cells can be distinguished from conventional memory cells by their low expression of CD49d (Haluszczak et al., 2009). Based on CD49d staining, the majority of expanded memory phenotype Dock2^*hsd/hsd*^ splenic T cells resemble virtual memory cells (Figure 1C). The ratio of virtual to true memory is also significantly increased in the absence of DOCK2 (Figure 1C).

### *Dock2^hsd/hsd^* virtual memory cells are functional and their presence correlates with protection from intracellular infection

A key feature of memory cells from antigen-inexperienced mice is the innate-like propensity for the rapid and cognate antigen-independent production of interferon-γ (IFN-γ) following intracellular bacterial infection in response to pro-inflammatory cytokines such as IL-12 and IL-18 or NKG2D ligands (Berg et al., 2002, 2003; Soudja et al., 2012; Chu et al., 2013). Virtual memory T cells are dependent upon IL-15 for their generation, and also require the continued presence of IL-15 in order to maintain the levels of effector molecules necessary for antigen-independent bystander protection (White et al., 2016). Consistent with these studies, a higher proportion of *Dock2^hsd/hsd^* CD8^+^ T cells respond rapidly to *in vitro* polyclonal stimuli (PMA + ionomycin, or IL-12 + IL-18) by secreting IFN-γ (Figure 1D and 1E).

As virtual memory cells can robustly traffic to the liver (White et al., 2016), we hypothesized that increased IFN-γ production by *Dock2*-deficient T cells could be protective against infection with *Listeria monocytogenes*, which replicates extensively in the liver. Indeed, expansion of virtual memory cells in the absence of DOCK2 correlated with increased resistance to *L. monocytogenes* infection, as *Dock2^hsd/hsd^* mice showed significantly lower bacterial burden 3 days after intravenous infection (Supplementary Figure 1). Importantly, there were no differences between wild type and *Dock2^hsd/hsd^* liver bacterial CFUs 18 hours post-infection, consistent with published studies showing that the protective effect of CD8^+^ derived IFN-γ was manifest only 3 days after infection (Berg et al., 2003).

### *Dock2^hsd/hsd^* memory phenotype T cells arise in a hematopoietic-intrinsic manner following thymic egress

Some studies have identified key roles for radio-resistant stromal cells in maintaining peripheral T cell homeostasis (Roozendaal and Mebius, 2011). With this in mind, we sought to evaluate the contribution of non-hematopoietic cells in the expansion of virtual memory T cells by transferring bone marrow into irradiated RAG-deficient recipients. We found that only recipients of Dock2^*hsd/hsd*^ bone marrow had a robust expansion of memory cells, while mice that received wild type bone marrow had a much smaller virtual memory compartment (Figure 2A). These experiments could not be performed in the setting of competitive reconstitution as *Dock2*-deficient hematopoietic progenitors exhibit a severe defect in bone marrow reconstitution under competition from wild-type cells due to an impaired response to CXCL12 (Kikuchi et al., 2008). Indeed, mixed bone marrow chimeras, where wild-type (WT) and *Dock2^hsd/hsd^* mice bone marrow were co-injected into the same *Rag1*^−/−^ recipient, resulted in ~20:1 hematopoietic reconstitution despite a 1:1 transfer of bone marrow precursors.

**Figure 2:**
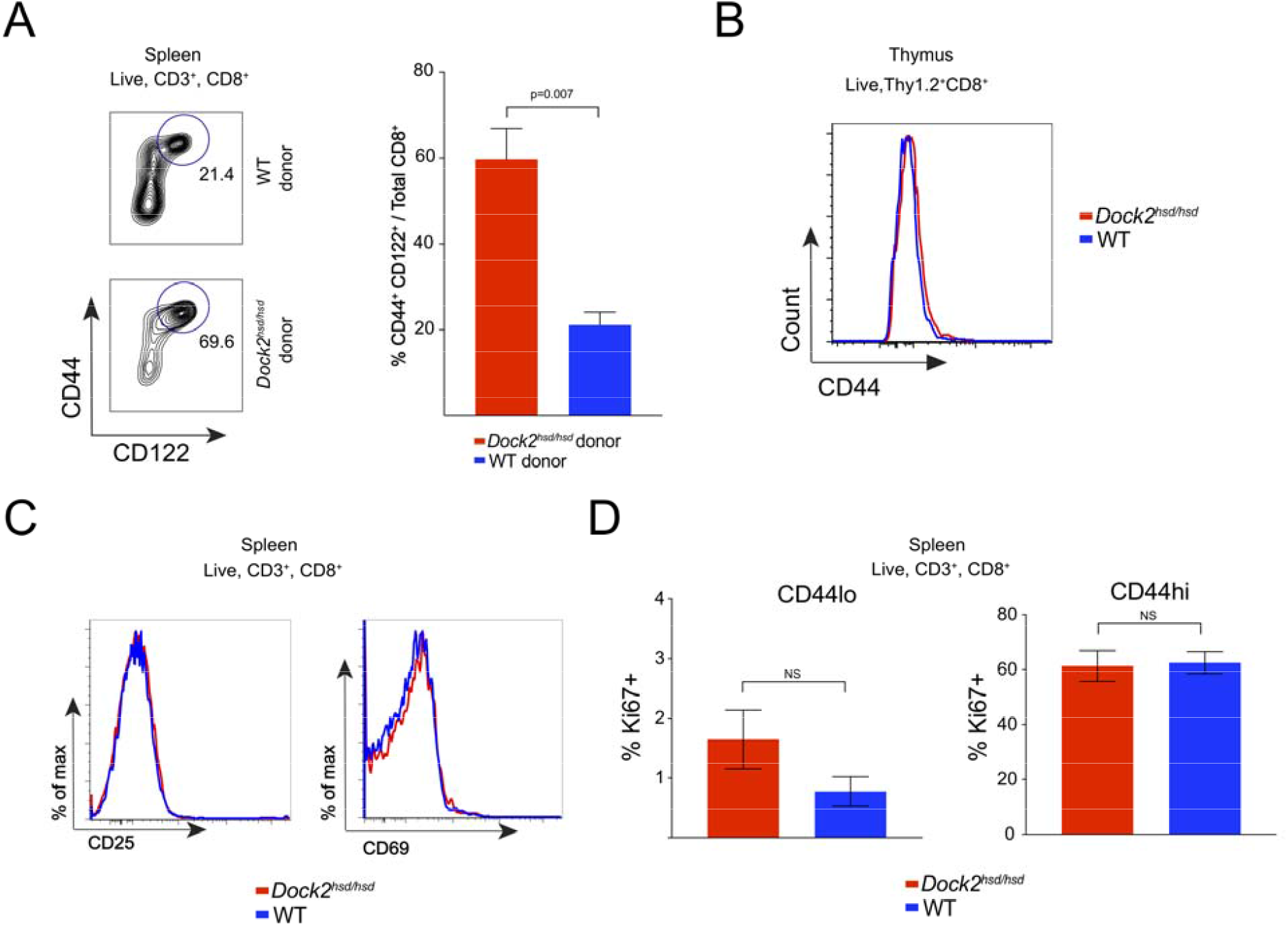
*Dock^hsd/hsd^* virtual memory T cells arise in a cell-intrinsic manner following thymic egress. (A) Bone marrow cells from either *Dock2^hsd/hsd^* or WT mice were injected into irradiated *Rag1*^−/−^ mice. The spontaneous generation of memory CD8^+^ T cells (CD44^hi^ CD122^hi^) was examined in the recipient mice at 12 weeks after cell transfer. (B) CD44 levels on the Thy1.2^+^ CD8^+^ CD4^−^ single positive thymocytes from WT and *Dock2*-deficient mice. (C) Levels of activation markers (CD25 and CD69) on splenic CD8^+^ T cells from WT and *Dock2*-deficient mice (D) Percentage of Ki-67+ cells in the CD44^lo^ and CD44^hi^ CD8^+^ T cell compartments in WT and *Dock2*-deficient mice. All experiments were performed twice in groups of 3 to 4 mice. Statistical significance was assessed using unpaired two-tailed Student’s t-test.

In contrast to innate CD8^+^ T cells that are generated prior to thymic egress (Berg, 2007; Lee et al., 2011), *Dock2^hsd/hsd^* virtual memory cells arise in the periphery, as mature CD8^+^CD4^−^CD44^hi^ cells are not present in the thymus (Figure 2B). Dock2^hsd/hsd^ deficient naive T cells also show no signs of early effector activation (surface CD69 and CD25 expression) or proliferation (as seen by Ki-67 expression) that accompany conventional true memory cell generation following thymic egress (Kaech and Cui, 2012) (Figure 2C & 2D).

### *Dock2^hsd/hsd^* naive T cells directly convert into memory phenotype cells

To explore the mechanism underlying the enhanced generation of *Dock2^hsd/hsd^* virtual memory cells, we compared the transcriptional profiles of *Dock2*-deficient and WT naive CD8^+^ T cells. Interestingly, we found that genes upregulated in *Dock2^hsd/hsd^* naive T cells are significantly enriched in several gene sets associated with memory T cell differentiation (Figure 3A). Consistent with the direct conversion of these mutant naïve CD8^+^ T cells to memory T cells and the bypassing of effector T cell clonal expansion, we observed no enrichment of any gene sets associated with early T cell activation.

**Figure 3:**
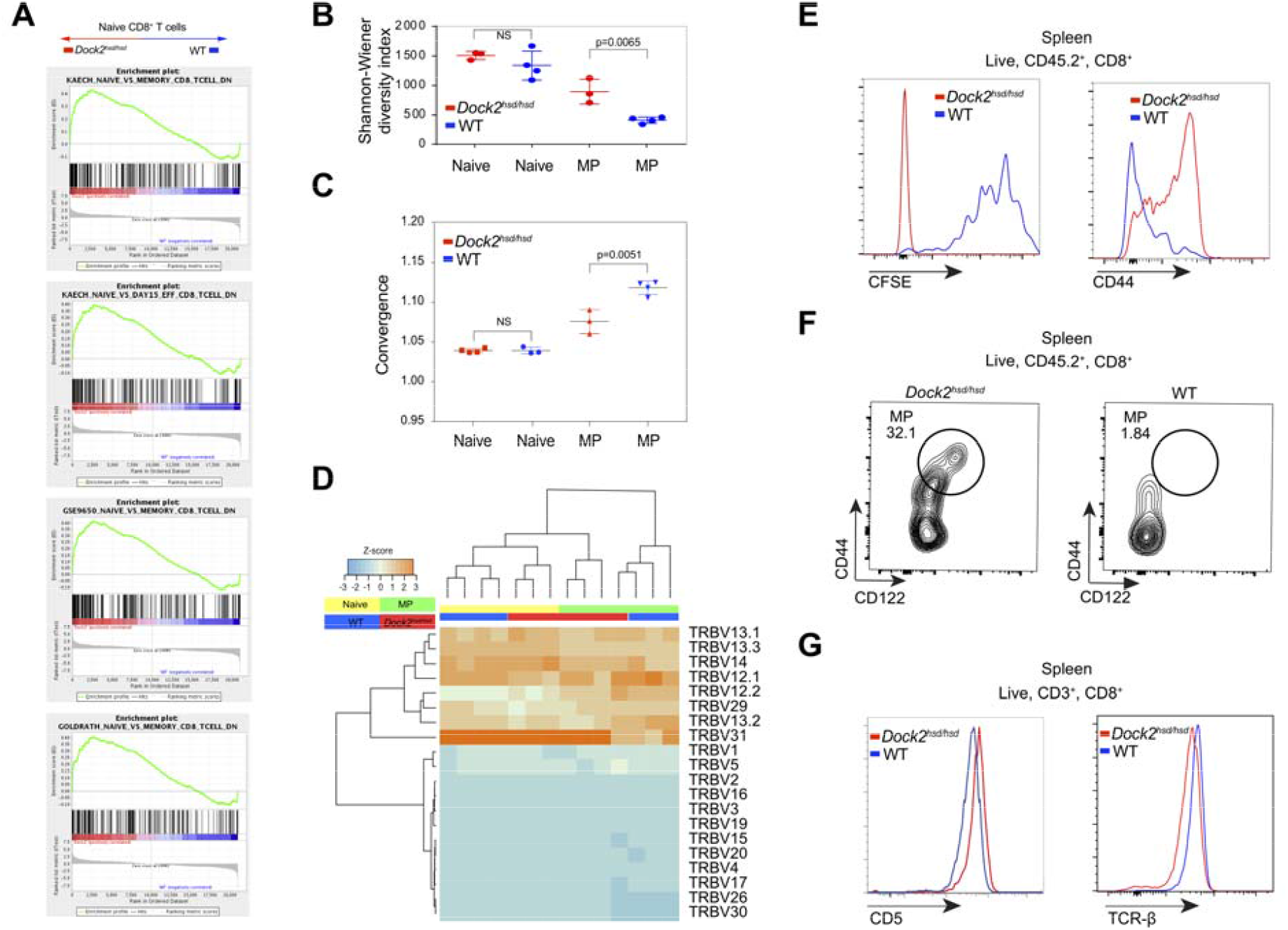
Naïve *Dock2^hsd/hsd^* T cells undergo spontaneous conversion into virtual memory T cells. (A) Gene set enrichment analysis of genes upregulated in CD8^+^ naive (CD44^lo^CD122^--^) *Dock2^hsd/hsd^* T cells compared to WT naive CD8^+^ T cells. (B and C) The Shannon-Weaver diversity index (B) and repertoire convergence (C) (i.e. number of unique CDR3 nucleotide sequences that encode the same amino acid sequence) in naive (CD44^lo^CD122^−^) and MP (CD44^hi^CD122^+^) T cells from WT and *Dock2^hsd/hsd^* mice. Statistical significance was assessed using unpaired two-tailed Student’s t test. (D) A hierarchically clustered heatmap of the frequency of TCR Vβ gene segment usage in naive (CD44^lo^CD122^--^) and MP (CD44^hi^CD122^+^) CD8^+^ T cells from *Dock^hsd/hsd^* and WT mice. (E) Naive T cells from CD45.2^+^ Thy1.2^+^ *Dock2^hsd/hsd^* and CD45.2^+^ Thy1.1^+^ WT mice were co-transferred into irradiated CD45.1^+^ Thy1.2^+^ lymphopenic mice and assessed for CFSE dilution and CD44 upregulation after 1 week. (F) Naive T cells from CD45.2^+^ Thy1.2^+^ *Dock2^hsd/hsd^* and CD45.2^+^ Thy1.1^+^ WT mice were co-transferred into unmanipulated lymphoreplete CD45.1^+^ Thy1.2^+^ mice and assessed for upregulation of memory markers after 3 weeks. (G) Surface levels of CD5 and TCR on CD8^+^ T cells from *Dock2^hsd/hsd^* and WT mice as assessed by flow cytometry.

TCR sequencing analysis suggested that the repertoire of the naive CD8^+^ T cells was comparably diverse in both *Dock2*-deficient and WT mice (Figure 3B). However, the *Dock2*-deficient virtual memory CD8^+^ cells were more diverse than WT virtual memory cells, suggesting that the conversion to virtual memory cells was a highly polyclonal process in *Dock2*-deficient mice. One measure of antigenic selection in a TCR repertoire is convergence, the number of unique CDR3 sequences encoding the same amino acid sequence (Shugay et al., 2015). In the context of virtual memory T cells, convergence can be interpreted as selection by self-antigens in the periphery. Analysis of the *Dock2^hsd/hsd^* memory T cell repertoire revealed a significantly lower number of unique CDR3 nucleotide sequences that encoded the same amino acid sequence, compared with wild type memory cells (Figure 3C). This decrease in convergence suggests that the self-antigen affinity threshold for entering the virtual memory compartment is lowered in the absence of DOCK2, allowing more “naive” T cells to enter this compartment. Further supporting this conclusion, we found that the TCR Vβ expression profile of *Dock2^hsd/hsd^* virtual memory cells bore remarkable similarity to naive T cells and is quite distinct from wild type virtual memory cells (Figure 3D).

In order to determine whether Dock2^*hsd/hsd*^ naive CD8^+^ T cells have an increased cell intrinsic propensity to convert to virtual memory, we investigated how adoptively cotransferred Dock2^*hsd/hsd*^ and WT naive T cells respond to homeostatic signals *in vivo. Dock2^hsd/hsd^* T cells transferred into irradiated lymphopenic mice proliferate at a strikingly faster rate than co-transferred WT T cells with concomitant CD44 upregulation (Figure 3E). Importantly, this increased rate of conversion to memory was also observed when naive *Dock2^hsd/hsd^* T cells were transferred into lymphoreplete mice without any irradiation (Figure 3F).

Surface CD5 levels, a proxy for tonic TCR signalling (Azzam et al., 1998, 2001), were significantly higher on naive T cells from Dock2^*hsd/hsd*^ mice than WT mice (Figure 3G). In addition, Dock2^*hsd/hsd*^ naive T cells exhibit lower surface TCR expression (Figure 3G). These findings suggest that an increase in self peptide-MHC triggering may be associated with the expansion of memory phenotype cells. Interestingly, increased CD5 levels are also seen in *Dock2^hsd/hsd^* CD8 single-positive thymocytes, suggesting that the *Dock2*-deficient cells may receive stronger positively selecting signals. However, this is not accompanied by dramatic changes in the proportions of thymic precursors (Supplementary Figure 2).

### Responsiveness to weak agonists is selectively enhanced in *Dock2*-deficient T cells

As our experiments implicated tonic TCR signaling in driving the conversion to virtual memory, we generated *Dock2*-deficient OT-I transgenic mice (OT-I *Dock2^hsd/hsd^*) to interrogate this phenomenon in the context of a defined TCR repertoire. OT-I *Dock2^hsd/hsd^* mice were born at Mendelian ratios and exhibited no obvious developmental defects. However, a close examination of T cells in the spleens of these mice revealed a striking accumulation of virtual memory cells, as compared to OT-I mice with normal expression of DOCK2 (Figure 4A, Supplementary Figure 3). Thus, the *Dock2*-dependent expansion of CD8^+^ T cells does not require a polyclonal repertoire.

**Figure 4:**
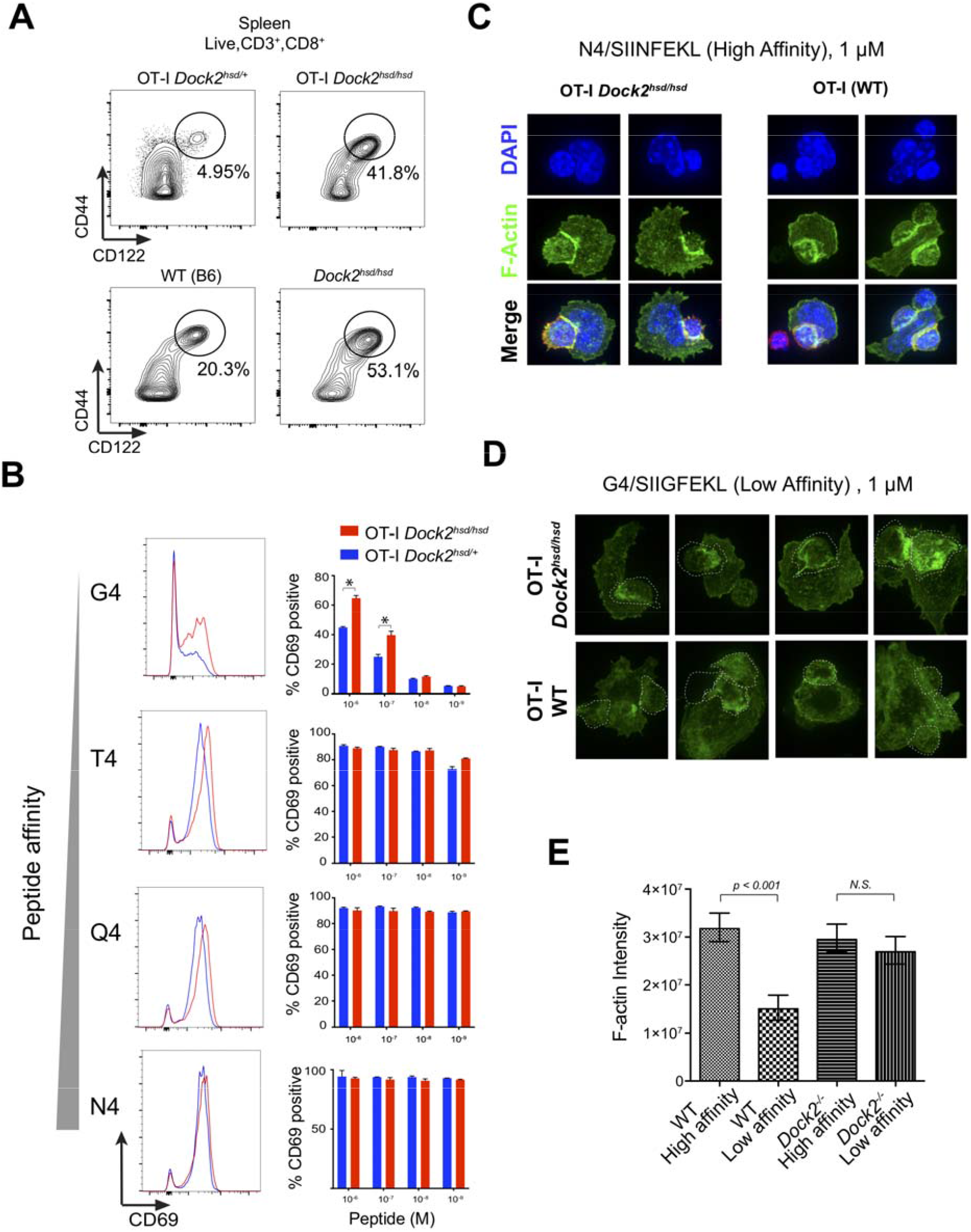
OT-I *Dock2^hsd/hsd^* mice have an expansion of memory phenotype (MP) T cells and show increased *ex vivo* responses to TCR stimulation. (A) The proportion of CD8^+^ CD44^hi^ CD122^+^ memory phenotype (MP) cells in the spleens of OT-I TCR-transgenic mice in the WT and *Dock2^hsd/hsd^* background. (B) CD69 upregulation in response to four hours of ex vivo stimulation with peptides of varying signal TCR affinity. Histograms of CD69 staining by flow cytometry show representative responses to stimulation with 1μM of peptide while bar graphs show the responses observed across a range of peptide concentrations. Statistically significant differences (p < 0.01) are marked by an asterisk. (C and D) Confocal microscopy of filamentous actin (green) at the immunological synapse in OT-I WT or OT-I *Dock2^hsd/hsd^* T cells co-cultured with syngeneic wild-type splenocytes in presence of (C) the high affinity SIINFEKL (N4) peptide or (D) the low affinity SIIGFEKL (G4) peptide. OT-I T cells were pre-stained with a fluorescently labeled anti-CD8 nanobody (red). (E) Quantification of F-actin intensity in OT-I WT and *Dock2^hsd/hsd^* CD8^+^ T cells upon stimulation with high (N4) or low (G4) affinity peptides shown as integrated fluorescence intensity. The Mann–Whitney unpaired t-test was used to assess statistical significance. The bars charts depict mean ± SEM.

We next measured the responses of naive *Dock2*-deficient OT-I T cells to altered peptide ligands with varying degrees of affinity for the OT-I TCR using the following peptides, listed in increasing order of affinity: SIIGFEKL (G4), SIITFEKL (T4), SIIQFEKL (Q4), and SIINFEKL (N4) (Salmond et al., 2014). We used CD69 upregulation (Cibrián and Sánchez-Madrid, 2017) and cytoskeletal remodeling (Kumari et al., 2015, 2014) to evaluate TCR-dependent responses. Incubation of splenocytes with strong and intermediate affinity agonists (N4, T4 and Q4) consistently activated OT-I T cells, regardless of DOCK2 expression (Figure 4B). However, *ex vivo* weak agonist (G4) stimulation resulted in significantly more CD69^+^ *Dock2*-deficient OT-I T cells when compared to cells from OT-I *Dock2^+/hsd^* littermates (Figure 4B).

TCR signaling is associated with actin polymerization and the enrichment of filamentous actin (F-actin) at the immunological synapse. Remodeling of F-actin at the immunological synapse is essential for TCR signaling events (Kumari et al., 2015, 2014). The high affinity (N4) peptide induced comparable levels of polymerized actin at the synapse in both *Dock2*-deficient as well as WT OT-I T cells (Figure 4C and 4E). However, in the *Dock2*-deficient OT-I T cells but not wild-type OT-I T cells, the lowest affinity peptide (G4) induced prominent actin polymerization (Figure 4D and 4E). Collectively, these findings suggest that *Dock2*-deficient OT-I T cells have a significantly reduced threshold of TCR activation in response to low affinity peptides.

## DISCUSSION

We had previously shown that the loss of DOCK2 results in a prominent expansion of CD8^+^ blood memory phenotype (MP) cells, and had used this phenotype to identify a loss-of-function *Dock2* genetic variant (*Dock2^hsd/hsd^*) in the commercially available C57BL6/NHsd substrain of C57BL/6 mice (Mahajan et al., 2016). In this study, we show that a similar expansion of CD8^+^ memory T cells is not specific to this *Dock2* mutant allele and is also seen in gene targeted *Dock2* knockout mice (Fukui et al., 2001). In contrast to conventional antigen-experienced memory cells, the majority of *Dock2^hsd/hsd^* memory T cells exhibit low surface expression of CD49d, originate in the periphery, lack activation makers, and have a highly diverse repertoire without any dominant clones. It can thus be concluded that the MP cells in *Dock2^hsd/hsd^* mice represent *bona fide* virtual memory cells.

Like innate CD8^+^ T lymphocytes, virtual memory cells can synthesize and secrete cytokines in an antigen-independent manner and in some cases, also provide bystander protection independent of TCR recognition. Despite prior reports of immune dysfunction in multiple hematopoietic lineages including lymphocytes in *Dock2*-deficient mice, we were surprised to find that *Dock2^hsd/hsd^* mice were more resistant to *Listeria monocytogenes* infection, likely due to the increased *ex vivo* innate-like IFN-γ production by T cells from these mice. Importantly, resistance to infection manifests itself 3 days post-infection and the kinetics of protection are consistent with innate-like immune protection by memory T cells; it has been shown that memory CD8^+^ T cells can impart protection against *Listeria monocytogenes* infection in an antigen-independent manner by producing high levels of IFN-γ in response to innate immune sources of IL-12 and IL-18 early in infection (Berg et al., 2003). It is also possible that the overlap between the replicative niche of this bacterium and extensive trafficking of virtual memory cells to the liver contributes to the protection (White et al., 2016). Although the loss of DOCK2 has been associated with immunodeficiency (Liu et al., 2016; Dobbs et al., 2015), we speculate that *Dock2* loss in mice and humans may partially protect from certain infections that overlap with anatomical sites preferred by virtual memory cells, but increase susceptibility to other pathogens that replicate at sites where virtual memory cells are present at a lower frequency.

Virtual memory T cell generation and maintenance is dependent on TCR triggering by self-peptide MHC and common γ-chain cytokine derived signals. However, there is little known about negative regulators of the virtual memory compartment (Fiege et al., 2015). In this study, we have provided evidence that DOCK2 sets a threshold for direct entry of naive CD8^+^ T cells into the virtual memory compartment. We have shown that there is reduced TCR sequence convergence and greater naive-like TCR Vβ usage in the *Dock2^hsd/hsd^* virtual memory TCR repertoire suggesting that more “naive” TCRs are able to enter this compartment. We found a striking enrichment in gene sets associated with memory differentiation in the genes that are upregulated in *Dock2^hsd/hsd^* naive CD8^+^ T cells relative to WT naive T cells. Notably, there was no enrichment of gene sets associated with effector differentiation. We observed an increased propensity of adoptively co-transferred *Dock2^hsd/hsd^* naive T cells to convert to a memory phenotype in response to homeostatic signals. We also demonstrated that *Dock2*-deficiency increases TCR sensitivity to weak agonists, and a higher level of CD5 observed on *Dock2^hsd/hsd^* T cells is consistent with enhanced responsiveness to self-peptide MHC *in vivo*.

A prior study by Sanui *et al*. showed that in the absence of DOCK2, MHC-II restricted 2B4 TCR transgenic CD4^+^ T cells exhibit a reduced response to weak agonist peptides (Sanui et al., 2003). However, the interpretation of this experiment may be complicated by the observation that DOCK2 may negatively influence lymphocyte proliferation independent of antigen receptor signaling, (Wang et al., 2010). Besides DOCK2, FYB1 has been shown to negatively regulate the size of the virtual memory compartment (Fiege et al., 2015). However, the relationship between DOCK2 and FYB1 has not yet been studied, and it is formally possible that DOCK2 and FYB1 function in the same or parallel pathways.

Recent studies have suggested that IL-4 and IL-15 derived from iNKT cells, control the size of the peripheral memory-like T cell compartment (Tripathi et al., 2016; Kurzweil et al., 2014; Akue et al., 2012). However, iNKT cell numbers are reduced in *Dock2^hsd/hsd^* mice (Mahajan et al. 2016) and the expansion of virtual memory cells in these mice is unlikely to result from changes in cytokine producing cells. Furthermore, an abundance of homeostatic cytokines cannot explain the selective expansion of virtual memory cells without impacting conventional memory T cells.

TCR sensitivity to self-peptides regulates virtual memory T cell formation (Kieper et al., 2004). Profiling reporters of tonic signalling, CD5 upregulation (Azzam et al., 2001) and TCR downregulation (Itoh et al., 1999) suggested that spontaneous conversion of *Dock2^hsd/hsd^* naive T cells to memory may be linked to increased TCR responsiveness to self-peptide MHC. We further probed this notion in the context of a restricted repertoire by generating *Dock2^hsd/hsd^* mice that carry the OT-I TCR transgene. Examination of OT-I *Dock2^hsd/hsd^* TCR transgenic mice revealed a robust expansion of OT-I T cells with memory markers in the absence of a polyclonal repertoire, exceeding what was observed in *Dock2^hsd/+^* littermates. We then directly assessed the consequence of DOCK2 loss on TCR triggering by using altered peptide ligands with varying levels of affinity to the OT-I TCR (Salmond et al., 2014). Interestingly, OT-I T cells were uniformly activated in response to strong (N4) and intermediate (Q4 and T4) peptide stimulation. However, stimulation with a weak agonist peptide (G4) resulted in a pronounced increase in the percentage of activated *OT-I Dock2^hsd/hsd^* T cells relative to OT-I T cells, consistent with our *in vivo* findings of increased self-peptide-MHC triggering. Therefore, we concluded that increased tonic TCR triggering likely drives direct conversion of naive T cells into virtual memory cells in *Dock2*-deficient mice. Triggering by self-peptide-MHC can also potentiate responses to common γ-chain cytokines (Palmer et al., 2011), and future studies will be needed to address the effect of DOCK2 loss on responses to cytokines.

It is currently unclear how the loss of DOCK2 sets the threshold for weak agonist TCR stimulation. One possibility is that the decreased interstitial motility (Nombela-Arrieta et al., 2007) observed by *Dock2*^−/−^ T cells results in increased TCR-MHC contact duration. It is possible that increases in antigen presenting cell “residency” time results in increased amounts of TCR triggering and concomitant conversion into memory cells. Another possibility is that the loss of DOCK2 disrupts the cortical network lowering the threshold for activation by weak ligands. Cortical actin forms a dense 100 nm thick layer lining the plasma membrane and can act as a barrier to TCR signaling by restricting the access of intracellular domains of LAT and CD3 to cytosolic signaling mediators such as PLC-γ1 in the absence of CD28 dependent costimulation (Dustin and Davis, 2014; Tan et al., 2014). The TCR and LAT are present in distinct microclusters in resting T cells, and the disruption of actin polymerization results in their activation-promoting aggregation (Lillemeier et al., 2010). DOCK2 is localized to the cell membrane via its interactions with the phospholipids, PIP3 and phosphatidic acid, and promotes RAC1-mediated actin polymerization (Fukui et al., 2001; Nishikimi et al., 2009, 2013). Therefore, we speculate that disruption of cortical actin in *Dock2*-deficient mice may contribute to the increased TCR responsiveness to weak agonists by promoting the enhanced diffusion of segregated transmembrane proteins such as the TCR and LAT and promote their association with downstream signaling mediators. Further studies on the role of Rac, the GTPase activated by DOCK2, in TCR signaling and virtual memory differentiation following egress into the periphery could also be informative as current studies of this GTPase have been, largely focused on thymic development and homing (Guo et al., 2008; Dumont et al., 2009).

In conclusion, we have demonstrated a novel role for DOCK2 in restricting the size of the virtual memory T cell compartment most likely by setting the threshold for responses against weak agonists. These findings suggest that any efforts to dampen immune responses using a small molecule inhibitor of DOCK2 should be tempered by an understanding that this protein has pleiotropic effects on peripheral T cell homeostasis (Nishikimi et al., 2012).

## MATERIALS AND METHODS

### Mice

*Dock2^hsd/hsd^* mice were purchased from Harlan Laboratories and maintained as a separate colony in a specific pathogen free environment in accordance with institutional guidelines. C57BL/6J, OT-I and RAG1 knockout mice were purchased from Jackson Laboratory. Unless otherwise specified, all experiments were conducted using 8-12 week-old mice.

### Bone marrow chimeras

Bone marrow was isolated from the indicated mice. Resuspended cells from the bone marrow were labeled using biotinylated anti-CD3 antibody and streptavidin microbeads (Miltenyi Biotec). Cells were then resuspended in PBS and 1×10^6^ cells were transferred to each mouse.

### Cell transfers

For lymphopenia induced proliferation experiments, mice were irradiated at a dose of 600 cGy. 6 hours later, a total of 1×10^6^ CFSE labeled naive T cells from *Dock2^hsd/hsd^* and WT mice were co-injected into irradiated congenic hosts. One week later, transferred cells were recovered and assessed for CFSE dilution and CD44 upregulation.

For experiments in lymphoreplete mice, a total of 2×10^6^ naive T cells from *Dock2^hsd/hsd^* and WT mice were co-injected into unmanipulated congenic hosts. Three weeks later, transferred cells were assessed for CD44 and CD122 upregulation.

### RNA Sequencing and TCR repertoire analysis

RNA was isolated with QIAGEN RNA isolation kits according to the manufacturer instructions. RNA-Seq libraries were then prepared using the Smart-Seq2 protocol (Picelli et al., 2013). Libraries were sequenced on an Illumina NextSeq 550. Paired end reads were aligned using the RSEM package and differentially expressed genes identified by EBSEQ with a PPDE cutoff of 0.95. Gene Set Enrichment Analysis (GSEA) was used to determine if the naïve CD8^+^ T cell gene expression program matched a known immunological gene expression signature (Subramanian et al., 2005; Godec et al., 2016). TCR repertoire libraries were prepared using the commercially available iRepertoire kit for mouse TCRβ and sequenced on a Illumina MiSeq instrument. V, D and J segment assignment and clonotypes identification was performed using MiXCR (Bolotin et al. 2015). VDJtools was used for post analysis determination of convergence and hierarchical clustering of samples based on TCR Vβ usage (Shugay et al., 2015).

### T cell stimulation

*Ex vivo* polyclonal stimulation was performed by incubating T cells in a cell stimulation cocktail from eBioscience (cat: 00-4975-03) for 4 hours. Cytokines were purchased from Peprotech and used at a concentration of 10 ng/ml for 18 hours. For OT-I TCR stimulation altered peptide ligands were synthesized by AnaSpec and used at the indicated concentrations to stimulate cells for 4 hours.

### Listeria infection

Mice were intravenously infected with 2×10^4^ CFUs of LM1043S. At the indicated times, livers were homogenized with 0.05% Triton-X in PBS followed by plating of serial dilutions on Brain Heart Infusion agar plates containing streptomycin.

### Flow cytometry

The following antibodies were used for surface staining CD8 (53-6.7), CD44 (IM7), CD49d (R1-2), CD122 (TM-β1), IFN-g (XMG1.2), CD69 (H1.2F3), CD25 (, Ki67 (16A8), TCR, Vβ5 (MR9-4), Vα2 (B20.1), CD45.1 (A20), CD45.2 (104), CD90.1 (OX-7), CD90.2 (30-H12). Cells were permeabilized for intracellular staining using the Foxp3 permeabilization kit (eBioscience cat:00-5523-00).

### Microscopy and image analysis

OT-I T cells tagged with fluorescently labeled anti-CD8 nanobodies were co-cultured for 4 hours in the presence of antigenic peptide-loaded syngeneic wild-type bone-marrow derived dendritic cells generated from B6 bone marrow (Rashidian et al., 2017; Inaba et al., 1992). The cells were fixed, stained and imaged using confocal microscopy as previously described (Kumari et al., 2015). Image processing was carried out using ImageJ and Metamorph software. To assess F-actin intensity, the region of interest was defined as corresponding to the zone of actin polymerization along the contact interface between T cell and dendritic cell, and the F-actin intensity was quantified using MetaMorph Software. All images presented here are raw images, displayed at identical contrast settings.

---

**Supplementary Figure 1:** Bacterial burden in the livers of *Listeria*-infected WT and *Dock2*-deficient mice at 18 and 72 hours post-infection. A representative flow cytometry plot is shown for each experiment to show the gating strategy. All experiments were performed twice with groups of 3 to 4 mice. Statistical significance was assessed using unpaired two-tailed Student’s t test.

**Supplementary Figure 2:** Flow cytometric analysis of the CD5 expression levels (left) on thymic T cell subsets and their respective proportions (right) in the thymi of WT and *Dock2^hsd/hsd^* mice.

**Supplementary Figure 3:** The proportion of CD8^+^ memory phenotype cells (MP) in the spleens of OT-I TCR transgenic mice in the Dock2^hsd/hsd^ and WT backgrounds, as well as Dock2^hsd/hsd^ and WT mice lacking the OT-I TCR transgene.

## REFERENCES

Akondy, R.S., M. Fitch, S. Edupuganti, S. Yang, H.T. Kissick, K.W. Li, B.A. Youngblood, H.A. Abdelsamed, D.J. McGuire, K.W. Cohen, G. Alexe, S. Nagar, M.M. McCausland, S. Gupta, P. Tata, W.N. Haining, M.J. McElrath, D. Zhang, B. Hu, W.J. Greenleaf, J.J. Goronzy, M.J. Mulligan, M. Hellerstein, and R. Ahmed. 2017. Origin and differentiation of human memory CD8 T cells after vaccination. Nature. doi:10.1038/nature24633.

Akue, A.D., J.-Y. Lee, and S.C. Jameson. 2012. Derivation and maintenance of virtual memory CD8 T cells. J. Immunol. 188:2516–2523.

Azzam, H.S., J.B. DeJarnette, K. Huang, R. Emmons, C.S. Park, C.L. Sommers, D. El-Khoury, E.W. Shores, and P.E. Love. 2001. Fine tuning of TCR signaling by CD5. J. Immunol. 166:5464–5472.

Azzam, H.S., A. Grinberg, K. Lui, H. Shen, E.W. Shores, and P.E. Love. 1998. CD5 expression is developmentally regulated by T cell receptor (TCR) signals and TCR avidity. J. Exp. Med. 188:2301–2311.

Berg, L.J. 2007. Signalling through TEC kinases regulates conventional versus innate CD8(+) T-cell development. Nat. Rev. Immunol. 7:479–485.

Berg, R.E., C.J. Cordes, and J. Forman. 2002. Contribution of CD8+ T cells to innate immunity: IFN-γ secretion induced by IL-12 and IL-18. Eur. J. Immunol. 32:2807–2816.

Berg, R.E., E. Crossley, S. Murray, and J. Forman. 2003. Memory CD8+ T cells provide innate immune protection against Listeria monocytogenes in the absence of cognate antigen. J. Exp. Med. 198:1583–1593.

Cheung, K.P., E. Yang, and A.W. Goldrath. 2009. Memory-like CD8+ T cells generated during homeostatic proliferation defer to antigen-experienced memory cells. J. Immunol. 183:3364–3372.

Chu, T., A.J. Tyznik, S. Roepke, A.M. Berkley, A. Woodward-Davis, L. Pattacini, M.J. Bevan, D. Zehn, and M. Prlic. 2013. Bystander-Activated Memory CD8 T Cells Control Early Pathogen Load in an Innate-like, NKG2D-Dependent Manner. Cell Rep. 3:701–708.

Cibrián, D., and F. Sánchez-Madrid. 2017. CD69: from activation marker to metabolic gatekeeper. Eur. J. Immunol. 47:946–953.

Dobbs, K., C. Domínguez Conde, S.-Y. Zhang, S. Parolini, M. Audry, J. Chou, E. Haapaniemi, S. Keles, I. Bilic, S. Okada, M.J. Massaad, S. Rounioja, A.M. Alwahadneh, N.K. Serwas, K. Capuder, E. Çiftçi, K. Felgentreff, T.K. Ohsumi, V. Pedergnana, B. Boisson, Ş. Haskoloğlu, A. Ensari, M. Schuster, A. Moretta, Y. Itan, O. Patrizi, F. Rozenberg, P. Lebon, J. Saarela, M. Knip, S. Petrovski, D.B. Goldstein, R.E. Parrott, B. Savas, A. Schambach, G. Tabellini, C. Bock, T.A. Chatila, A.M. Comeau, R.S. Geha, L. Abel, R.H. Buckley, A. Ikincioğullari, W. Al-Herz, M. Helminen, F. Doğu, J.-L. Casanova, K. Boztuğ, and L.D. Notarangelo. 2015. Inherited DOCK2 Deficiency in Patients with Early-Onset Invasive Infections. N. Engl. J. Med. 372:2409–2422.

Drobek, A., A. Moudra, D. Mueller, M. Huranova, V. Horkova, M. Pribikova, R. Ivanek, S. Oberle, D. Zehn, K.D. McCoy, P. Draber, and O. Stepanek. 2018. Strong homeostatic TCR signals induce formation of self-tolerant virtual memory CD8 T cells. EMBO J. 37. doi:10.15252/embj.201798518.

Dumont, C., A. Corsoni-Tadrzak, S. Ruf, J. de Boer, A. Williams, M. Turner, D. Kioussis, and V.L.J. Tybulewicz. 2009. Rac GTPases play critical roles in early T-cell development. Blood. 113:3990–3998.

Dustin, M.L., and S.J. Davis. 2014. TCR signaling: the barrier within. Nat. Immunol. 15:136–137.

Ernst, B., D.S. Lee, J.M. Chang, J. Sprent, and C.D. Surh. 1999. The peptide ligands mediating positive selection in the thymus control T cell survival and homeostatic proliferation in the periphery. Immunity. 11:173–181.

Fiege, J.K., B.J. Burbach, and Y. Shimizu. 2015. Negative Regulation of Memory Phenotype CD8 T Cell Conversion by Adhesion and Degranulation-Promoting Adapter Protein. J. Immunol. 195:3119–3128.

Fukui, Y., O. Hashimoto, T. Sanui, T. Oono, H. Koga, M. Abe, A. Inayoshi, M. Noda, M. Oike, T. Shirai, and T. Sasazuki. 2001. Haematopoietic cell-specific CDM family protein DOCK2 is essential for lymphocyte migration. Nature. 412:826–831.

Ge, Q., V.P. Rao, B.K. Cho, H.N. Eisen, and J. Chen. 2001. Dependence of lymphopenia-induced T cell proliferation on the abundance of peptide/ MHC epitopes and strength of their interaction with T cell receptors. Proc. Natl. Acad. Sci. U. S. A. 98:1728–1733.

Godec, J., Y. Tan, A. Liberzon, P. Tamayo, S. Bhattacharya, A.J. Butte, J.P. Mesirov, and W.N. Haining. 2016. Compendium of Immune Signatures Identifies Conserved and Species-Specific Biology in Response to Inflammation. Immunity. 44:194–206.

Goldrath, A.W., and M.J. Bevan. 1999. Low-affinity ligands for the TCR drive proliferation of mature CD8+ T cells in lymphopenic hosts. Immunity. 11:183–190.

Goldrath, A.W., C.J. Luckey, R. Park, C. Benoist, and D. Mathis. 2004. The molecular program induced in T cells undergoing homeostatic proliferation. Proc. Natl. Acad. Sci. U. S. A. 101:16885–16890.

Gotoh, K., Y. Tanaka, A. Nishikimi, A. Inayoshi, M. Enjoji, R. Takayanagi, T. Sasazuki, and Y. Fukui. 2008. Differential requirement for DOCK2 in migration of plasmacytoid dendritic cells versus myeloid dendritic cells. Blood. 111:2973–2976.

Gotoh, K., Y. Tanaka, A. Nishikimi, R. Nakamura, H. Yamada, N. Maeda, T. Ishikawa, K. Hoshino, T. Uruno, Q. Cao, S. Higashi, Y. Kawaguchi, M. Enjoji, R. Takayanagi, T. Kaisho, Y. Yoshikai, and Y. Fukui. 2010. Selective control of type I IFN induction by the Rac activator DOCK2 during TLR-mediated plasmacytoid dendritic cell activation. J. Exp. Med. 207:721–730.

Guo, F., J.A. Cancelas, D. Hildeman, D.A. Williams, and Y. Zheng. 2008. Rac GTPase isoforms Rac1 and Rac2 play a redundant and crucial role in T-cell development. Blood. 112:1767–1775.

Haluszczak, C., A.D. Akue, S.E. Hamilton, L.D.S. Johnson, L. Pujanauski, L. Teodorovic, S.C. Jameson, and R.M. Kedl. 2009. The antigen-specific CD8+ T cell repertoire in unimmunized mice includes memory phenotype cells bearing markers of homeostatic expansion. J. Exp. Med. 206:435–448.

Hamilton, S.E., and S.C. Jameson. 2008. The nature of the lymphopenic environment dictates protective function of homeostatic-memory CD8+ T cells. Proceedings of the National Academy of Sciences. 105:18484–18489.

Inaba, K., M. Inaba, N. Romani, H. Aya, M. Deguchi, S. Ikehara, S. Muramatsu, and R.M. Steinman. 1992. Generation of large numbers of dendritic cells from mouse bone marrow cultures supplemented with granulocyte/macrophage colony-stimulating factor. J. Exp. Med. 176:1693–1702.

Itoh, Y., B. Hemmer, R. Martin, and R.N. Germain. 1999. Serial TCR engagement and down-modulation by peptide:MHC molecule ligands: relationship to the quality of individual TCR signaling events. J. Immunol. 162:2073–2080.

Jacomet, F., E. Cayssials, S. Basbous, A. Levescot, N. Piccirilli, D. Desmier, A. Robin, A. Barra, C. Giraud, F. Guilhot, L. Roy, A. Herbelin, and J.-M. Gombert. 2015. Evidence for eomesodermin-expressing innate-like CD8(+) KIR/NKG2A(+) T cells in human adults and cord blood samples. Eur. J. Immunol. 45:1926–1933.

Jameson, S.C. 2002. Maintaining the norm: T-cell homeostasis. Nat. Rev. Immunol. 2:547–556.

Kaech, S.M., and W. Cui. 2012. Transcriptional control of effector and memory CD8+ T cell differentiation. Nat. Rev. Immunol. 12:749–761.

Kaech, S.M., J.T. Tan, E.J. Wherry, B.T. Konieczny, C.D. Surh, and R. Ahmed. 2003. Selective expression of the interleukin 7 receptor identifies effector CD8 T cells that give rise to long-lived memory cells. Nat. Immunol. 4:1191–1198.

Kieper, W.C., J.T. Burghardt, and C.D. Surh. 2004. A role for TCR affinity in regulating naive T cell homeostasis. J. Immunol. 172:40–44.

Kikuchi, T., S. Kubonishi, M. Shibakura, N. Namba, T. Matsui, Y. Fukui, M. Tanimoto, and Y. Katayama. 2008. Dock2 participates in bone marrow lympho-hematopoiesis. Biochem. Biophys. Res. Commun. 367:90–96.

Kim, J., D.-Y. Chang, H.W. Lee, H. Lee, J.H. Kim, P.S. Sung, K.H. Kim, S.-H. Hong, W. Kang, J. Lee, S.Y. Shin, H.T. Yu, S. You, Y.S. Choi, I. Oh, D.H. Lee, D.H. Lee, M.K. Jung, K.-S. Suh, S. Hwang, W. Kim, S.-H. Park, H.J. Kim, and E.-C. Shin. 2017. Innate-like Cytotoxic Function of Bystander-Activated CD8+ T Cells Is Associated with Liver Injury in Acute Hepatitis A. Immunity. doi:10.1016/j.immuni.2017.11.025.

Kumari, S., S. Curado, V. Mayya, and M.L. Dustin. 2014. T cell antigen receptor activation and actin cytoskeleton remodeling. Biochim. Biophys. Acta. 1838:546–556.

Kumari, S., D. Depoil, R. Martinelli, E. Judokusumo, G. Carmona, F.B. Gertler, L.C. Kam, C.V. Carman, J.K. Burkhardt, D.J. Irvine, and M.L. Dustin. 2015. Actin foci facilitate activation of the phospholipase C-γ in primary T lymphocytes via the WASP pathway. Elife. 4. doi:10.7554/eLife.04953.

Kurzweil, V., A. LaRoche, and P.M. Oliver. 2014. Increased peripheral IL-4 leads to an expanded virtual memory CD8+ population. J. Immunol. 192:5643–5651.

Lee, Y.J., S.C. Jameson, and K.A. Hogquist. 2011. Alternative memory in the CD8 T cell lineage. Trends Immunol. 32:50–56.

Le Floc’h, A., Y. Tanaka, N.S. Bantilan, G. Voisinne, G. Altan-Bonnet, Y. Fukui, and M. Huse. 2013. Annular PIP3 accumulation controls actin architecture and modulates cytotoxicity at the immunological synapse. J. Exp. Med. 210:2721–2737.

Lillemeier, B.F., M.A. Mörtelmaier, M.B. Forstner, J.B. Huppa, J.T. Groves, and M.M. Davis. 2010. TCR and Lat are expressed on separate protein islands on T cell membranes and concatenate during activation. Nat. Immunol. 11:90–96.

Liu, Z., S.M. Man, Q. Zhu, P. Vogel, S. Frase, Y. Fukui, and T.-D. Kanneganti. 2016. DOCK2 confers immunity and intestinal colonization resistance to Citrobacter rodentium infection. Sci. Rep. 6:27814.

Mahajan, V.S., E. Demissie, H. Mattoo, V. Viswanadham, A. Varki, R. Morris, and S. Pillai. 2016. Striking Immune Phenotypes in Gene-Targeted Mice Are Driven by a Copy-Number Variant Originating from a Commercially Available C57BL/6 Strain. Cell Rep. 15:1901–1909.

Min, B., R. McHugh, G.D. Sempowski, C. Mackall, G. Foucras, and W.E. Paul. 2003. Neonates support lymphopenia-induced proliferation. Immunity. 18:131–140.

Nishikimi, A., H. Fukuhara, W. Su, T. Hongu, S. Takasuga, H. Mihara, Q. Cao, F. Sanematsu, M. Kanai, H. Hasegawa, Y. Tanaka, M. Shibasaki, Y. Kanaho, T. Sasaki, M.A. Frohman, and Y. Fukui. 2009. Sequential regulation of DOCK2 dynamics by two phospholipids during neutrophil chemotaxis. Science. 324:384–387.

Nishikimi, A., M. Kukimoto-Niino, S. Yokoyama, and Y. Fukui. 2013. Immune regulatory functions of DOCK family proteins in health and disease. Exp. Cell Res. 319:2343–2349.

Nishikimi, A., T. Uruno, X. Duan, Q. Cao, Y. Okamura, T. Saitoh, N. Saito, S. Sakaoka, Y. Du, A. Suenaga, M. Kukimoto-Niino, K. Miyano, K. Gotoh, T. Okabe, F. Sanematsu, Y. Tanaka, H. Sumimoto, T. Honma, S. Yokoyama, T. Nagano, D. Kohda, M. Kanai, and Y. Fukui. 2012. Blockade of inflammatory responses by a small-molecule inhibitor of the Rac activator DOCK2. Chem. Biol. 19:488–497.

Nombela-Arrieta, C., T.R. Mempel, S.F. Soriano, I. Mazo, M.P. Wymann, E. Hirsch, C. Martínez-A, Y. Fukui, U.H. von Andrian, and J.V. Stein. 2007. A central role for DOCK2 during interstitial lymphocyte motility and sphingosine-1-phosphate-mediated egress. J. Exp. Med. 204:497–510.

Palmer, M.J., V.S. Mahajan, J. Chen, D.J. Irvine, and D.A. Lauffenburger. 2011. Signaling thresholds govern heterogeneity in IL-7-receptor-mediated responses of naïve CD8(+) T cells. Immunol. Cell Biol. 89:581–594.

Picelli, S., Å.K. Björklund, O.R. Faridani, S. Sagasser, G. Winberg, and R. Sandberg. 2013. Smart-seq2 for sensitive full-length transcriptome profiling in single cells. Nat. Methods. 10:1096–1098.

Rashidian, M., J.R. Ingram, M. Dougan, A. Dongre, K.A. Whang, C. LeGall, J.J. Cragnolini, B. Bierie, M. Gostissa, J. Gorman, G.M. Grotenbreg, A. Bhan, R.A. Weinberg, and H.L. Ploegh. 2017. Predicting the response to CTLA-4 blockade by longitudinal noninvasive monitoring of CD8 T cells. J. Exp. Med. 214:2243–2255.

Roozendaal, R., and R.E. Mebius. 2011. Stromal cell-immune cell interactions. Annu. Rev. Immunol. 29:23–43.

Salmond, R.J., R.J. Brownlie, V.L. Morrison, and R. Zamoyska. 2014. The tyrosine phosphatase PTPN22 discriminates weak self peptides from strong agonist TCR signals. Nat. Immunol. 15:875–883.

Sanui, T., A. Inayoshi, M. Noda, E. Iwata, M. Oike, T. Sasazuki, and Y. Fukui. 2003. DOCK2 is essential for antigen-induced translocation of TCR and lipid rafts, but not PKC-theta and LFA-1, in T cells. Immunity. 19:119–129.

Shugay, M., D.V. Bagaev, M.A. Turchaninova, D.A. Bolotin, O.V. Britanova, E.V. Putintseva, M.V. Pogorelyy, V.I. Nazarov, I.V. Zvyagin, V.I. Kirgizova, K.I. Kirgizov, E.V. Skorobogatova, and D.M. Chudakov. 2015. VDJtools: Unifying Post-analysis of T Cell Receptor Repertoires. PLoS Comput. Biol. 11:e1004503.

Smith, N.L., R.K. Patel, A. Reynaldi, J.K. Grenier, J. Wang, N.B. Watson, K. Nzingha, K.J. Yee Mon, S.A. Peng, A. Grimson, M.P. Davenport, and B.D. Rudd. 2018. Developmental Origin Governs CD8+ T Cell Fate Decisions during Infection. Cell. 174:117–130.e14.

Sosinowski, T., J.T. White, E.W. Cross, C. Haluszczak, P. Marrack, L. Gapin, and R.M. Kedl. 2013. CD8α+ dendritic cell trans presentation of IL-15 to naive CD8+ T cells produces antigen-inexperienced T cells in the periphery with memory phenotype and function. J. Immunol. 190:1936–1947.

Soudja, S.M., A.L. Ruiz, J.C. Marie, and G. Lauvau. 2012. Inflammatory monocytes activate memory CD8(+) T and innate NK lymphocytes independent of cognate antigen during microbial pathogen invasion. Immunity. 37:549–562.

Sprent, J., and C.D. Surh. 2011. Normal T cell homeostasis: the conversion of naive cells into memory-phenotype cells. Nat. Immunol. 12:478–484.

Subramanian, A., P. Tamayo, V.K. Mootha, S. Mukherjee, B.L. Ebert, M.A. Gillette, A. Paulovich, S.L. Pomeroy, T.R. Golub, E.S. Lander, and J.P. Mesirov. 2005. Gene set enrichment analysis: a knowledge-based approach for interpreting genome-wide expression profiles. Proc. Natl. Acad. Sci. U. S. A. 102:15545–15550.

Tan, J.T., E. Dudl, E. LeRoy, R. Murray, J. Sprent, K.I. Weinberg, and C.D. Surh. 2001. IL-7 is critical for homeostatic proliferation and survival of naive T cells. Proc. Natl. Acad. Sci. U. S. A. 98:8732–8737.

Tan, Y.X., B.N. Manz, T.S. Freedman, C. Zhang, K.M. Shokat, and A. Weiss. 2014. Inhibition of the kinase Csk in thymocytes reveals a requirement for actin remodeling in the initiation of full TCR signaling. Nat. Immunol. 15:186–194.

Terasawa, M., T. Uruno, S. Mori, M. Kukimoto-Niino, A. Nishikimi, F. Sanematsu, Y. Tanaka, S. Yokoyama, and Y. Fukui. 2012. Dimerization of DOCK2 is essential for DOCK2-mediated Rac activation and lymphocyte migration. PLoS One. 7:e46277.

Tripathi, P., S.C. Morris, C. Perkins, A. Sholl, F.D. Finkelman, and D.A. Hildeman. 2016. IL-4 and IL-15 promotion of virtual memory CD8+ T cells is determined by genetic background. Eur. J. Immunol. 46:2333–2339.

Van Kaer, L. 2015. Innate and virtual memory T cells in man. Eur. J. Immunol. 45:1916–1920.

Wang, L., H. Nishihara, T. Kimura, Y. Kato, M. Tanino, M. Nishio, M. Obara, T. Endo, T. Koike, and S. Tanaka. 2010. DOCK2 regulates cell proliferation through Rac and ERK activation in B cell lymphoma. Biochem. Biophys. Res. Commun. 395:111–115.

White, J.T., E.W. Cross, M.A. Burchill, T. Danhorn, M.D. McCarter, H.R. Rosen, B. O’Connor, and R.M. Kedl. 2016. Virtual memory T cells develop and mediate bystander protective immunity in an IL-15-dependent manner. Nat. Commun. 7:11291.

White, J.T., E.W. Cross, and R.M. Kedl. 2017. Antigen-inexperienced memory CD8(+) T cells: where they come from and why we need them. Nat. Rev. Immunol. doi:10.1038/nri.2017.34.

Yu, B., K. Zhang, J. Justin Milner, C. Toma, R. Chen, J.P. Scott-Browne, R.M. Pereira, S. Crotty, J.T. Chang, M.E. Pipkin, W. Wang, and A.W. Goldrath. 2017. Epigenetic landscapes reveal transcription factors that regulate CD8+ T cell differentiation. Nat. Immunol. 18:573–582.

Zhang, X., B. Mozeleski, S. Lemoine, E. Dériaud, A. Lim, D. Zhivaki, E. Azria, C. Le Ray, G. Roguet, O. Launay, A. Vanet, C. Leclerc, and R. Lo-Man. 2014. CD4 T cells with effector memory phenotype and function develop in the sterile environment of the fetus. Sci. Transl. Med. 6:238ra72.

