## Supplementary figures and images for "DOCK2 sets the threshold for entry into the virtual memory CD8^+^ T cell compartment by negatively regulating tonic TCR triggering"

### Supplementary Figure 1

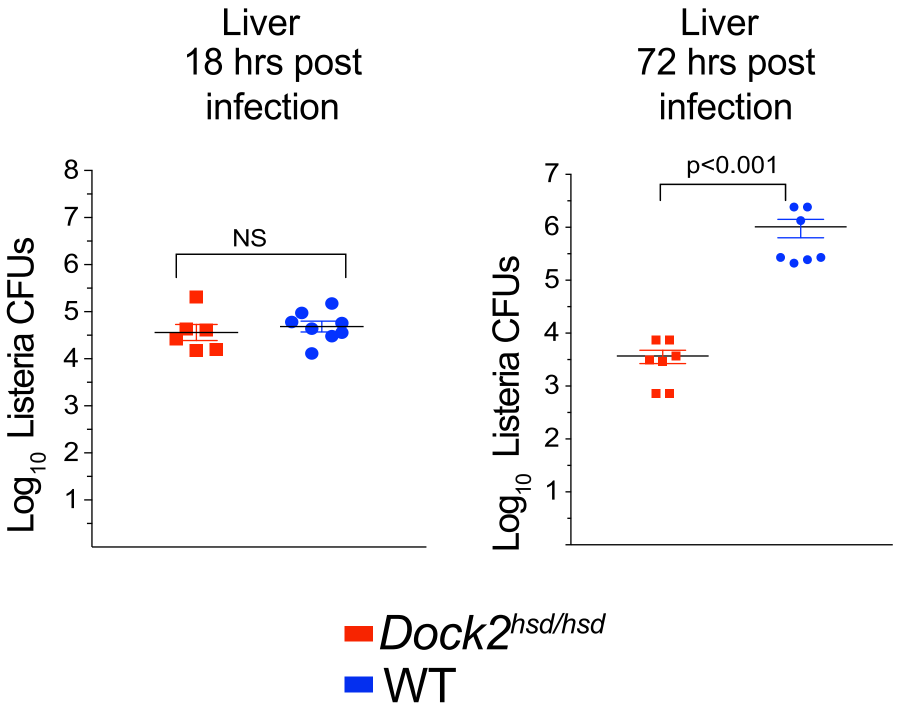

### Supplementary Figure 2

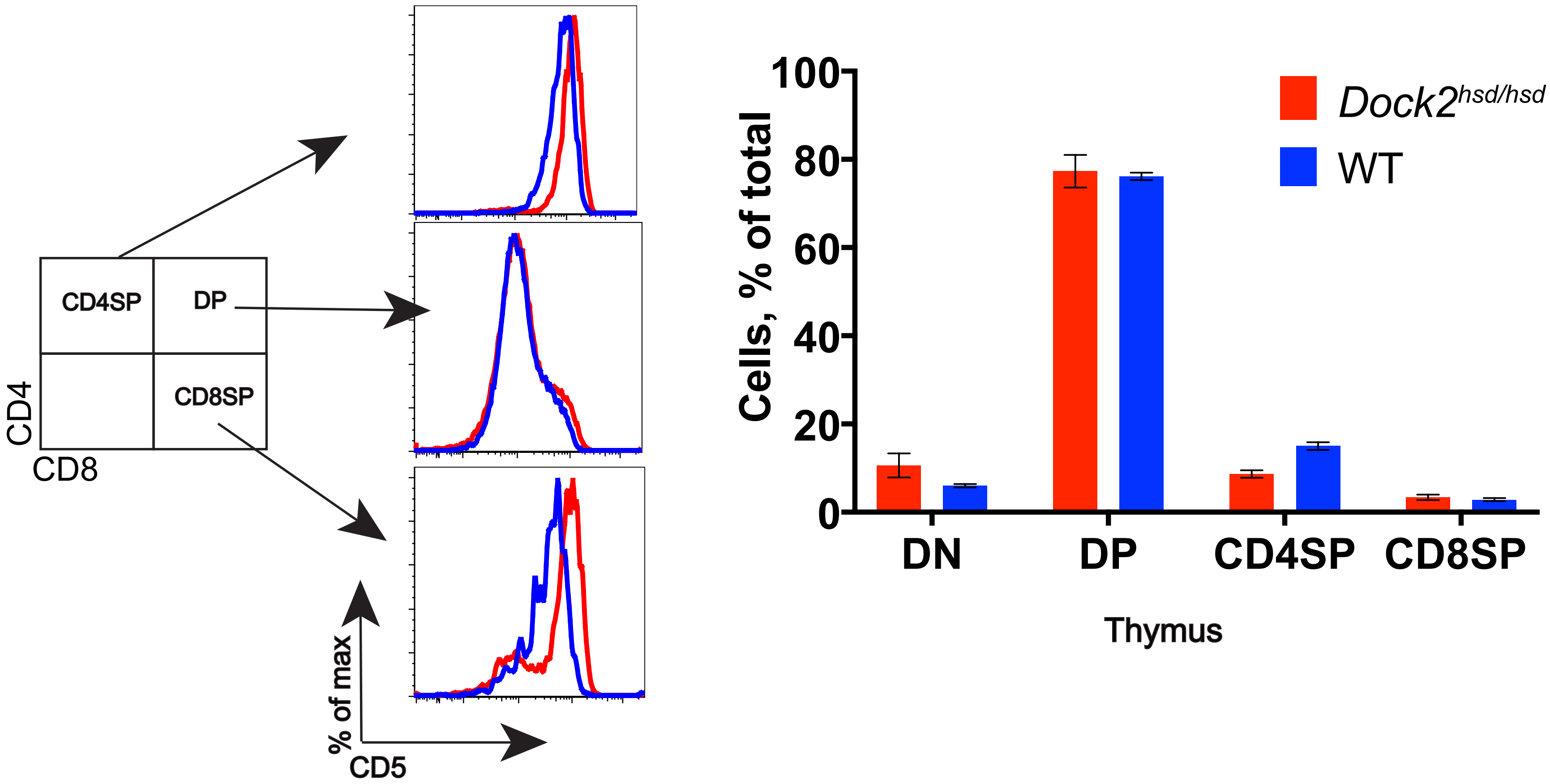

### Supplementary Figure 3

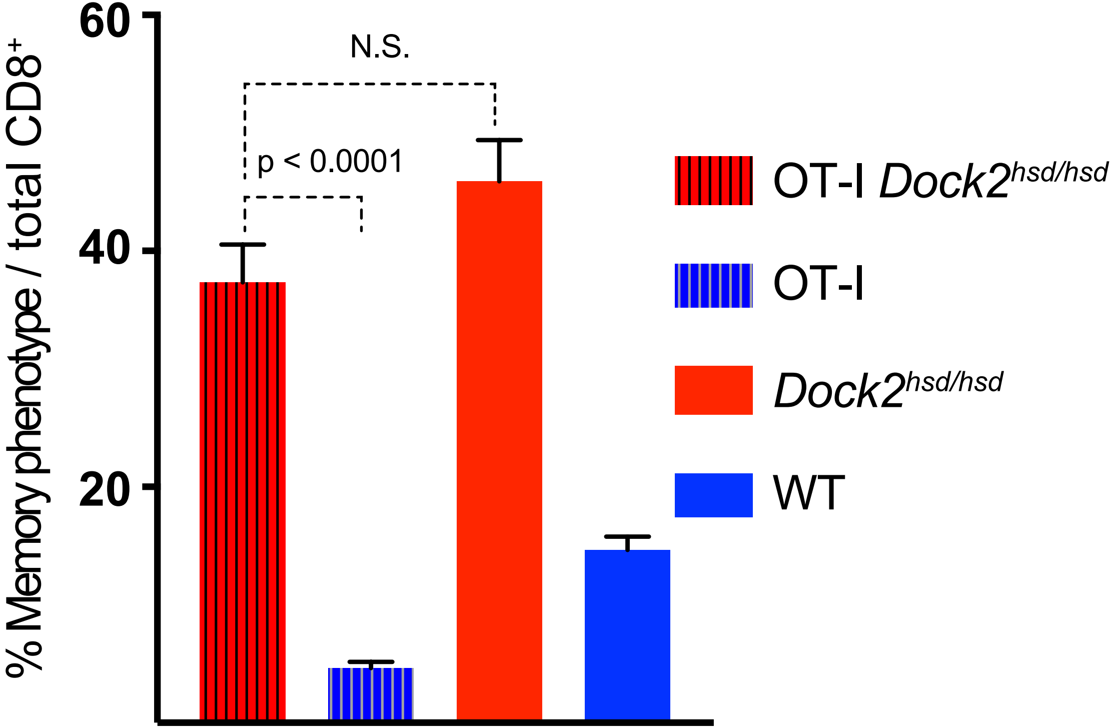
